# A Kidney Stone–Associated CLDN4 Variant Impairs Tight Junction Stability and Paracellular Ion Permeability

**DOI:** 10.64898/2026.02.10.705174

**Authors:** Forough Chelangarimiyandoab, Manu Rajinder Kumar, Kristina McNaughton, Grace Essuman, Daniel G. Fuster, R. Todd Alexander, Emmanuelle Cordat

## Abstract

Claudin-4 (CLDN4) is a key determinant of paracellular ion transport in the distal nephron, where it contributes to chloride permeability and transepithelial resistance. Although CLDN4 knockout mice exhibit hypercalciuria, the epithelial mechanism linking CLDN4 to calcium permeability and kidney stone disease remains unclear. We examined the molecular and functional effects of a kidney stone–associated CLDN4 variant P74L which was identified in two unrelated individuals with nephrolithiasis from the Bern Kidney Stone Registry. Using doxycycline-inducible epithelial cell models expressing human wild-type (WT) or mutant CLDN4, we show that the P74L variant displayed reduced protein stability, impaired junctional incorporation, and decreased surface expression. In contrast to WT CLDN4, whose overexpression increased transepithelial electrical resistance and restricted paracellular sodium, chloride, and calcium permeability, P74L CLDN4 failed to confer these effects. Expression of P74L CLDN4 was associated with reduced CLDN3 and CLDN7 messenger abundance without significant changes in CLDN8 or transcriptional regulation of other distal calcium (and other ion) transport genes. Together, these findings identify CLDN4 P74L as a loss-of-function variant that increases epithelial calcium permeability, possibly leading to increased calcium back-flux in the distal nephron relevant to nephrolithiasis.

## INTRODUCTION

Kidney stones (KS) are a prevalent and recurrent disorder, with a lifetime incidence of up to 10% of the population, imposing a significant health and economic burden worldwide (1–3). Most KS are composed of calcium salts, primarily calcium oxalate or calcium phosphate, with hypercalciuria representing the principal risk factor for their formation (2). Hypercalciuria can arise from abnormalities that disrupt calcium handling, including genetic defects affecting the renal tubular transport or epithelial barrier integrity (4). The renal distal nephron plays a critical role in the fine regulation of calcium, sodium, chloride, and proton handling (2,4–9). These processes rely on coordinated transcellular and paracellular transport mechanisms that maintain ionic balance and transepithelial voltage (10).

Tight junctions (TJs) contribute to the paracellular barrier of epithelial tissues, maintaining polarity while regulating ion selectivity and solute permeability (11–14). The molecular composition of TJs in the distal nephron confers specific permeability properties that complement active transcellular transport processes (13,15–18). Claudins (CLDNs) are the principal determinants of TJ’s selective permeability (19). Disruption of claudin expression or function compromises epithelial integrity and ionic homeostasis, contributing to distal tubular dysfunction (20–29).

Claudin-4 (CLDN4) is abundantly expressed in the distal nephron, particularly the connecting tubule (CNT) and the collecting duct (CD), where it forms a chloride-permeable, cation-restrictive barrier in coordination with Claudin-8 (18,20,21). Functional and genetic studies in epithelial cells and knockout mouse models show that CLDN4 determines paracellular chloride permeability and thereby contributes to the transepithelial voltage that results from the electrogenic sodium reabsorption through the epithelial sodium channel (ENaC) (21,29). Loss or mislocalization of CLDN4 disrupts chloride conductance and epithelial polarity, resulting in impaired salt reabsorption and hypotension (21,25–28). In Madin–Darby canine kidney (MDCK) epithelial monolayers, CLDN4 can also modulate paracellular selectivity indirectly through “inter-claudin interference,” whereby it suppresses the channel activity of pore-forming claudins such as CLDN2 or CLDN15 without uniformly tightening the barrier (30). Therefore, CLDN4 plays a dual role as both a pore-forming and regulatory component of the epithelial barrier in the distal nephron.

In the distal nephron, calcium is reabsorbed via a transcellular pathway involving apical uptake through the transient receptor potential vanilloid 5 (TRPV5) channel, cytosolic buffering by Calbindin D28K (also known as Calb28k or CaBP28k), and basolateral extrusion mediated by the Na⁺/Ca²⁺ exchanger 1 (NCX1) and plasma-membrane Ca²⁺-ATPase (PMCA) isoforms PMCA4 and PMCA1 (31–34). These processes occur within an epithelium where electrogenic Na⁺ reabsorption via ENaC generates a lumen-negative transepithelial voltage (35). Under these conditions, increased paracellular cation permeability would be predicted to favor electrochemically driven calcium back-flux from the interstitium into the tubular lumen, counteracting net calcium reabsorption.

Although *Cldn4* knockout mice exhibit increased fractional excretion of calcium compared with littermates (28), the role of CLDN4 in calcium homeostasis and kidney stone formation remains unclear. Whole-exome sequencing from the Bern Kidney Stone Registry (BKSR) identified *CLDN4* variants in two unrelated male patients with recurrent calcium nephrolithiasis and osteopenia (36). These individuals carried a proline-to-leucine substitution at position 74 (CLDN4 P74L) within the highly conserved first extracellular loop (ECL1), a region that determines paracellular ion selectivity among claudin family members (29,37–40). Given its location adjacent to the predicted pore-lining region, the P74L variant is predicted to alter TJ structure and ionic selectivity (21,37–40). Therefore, we hypothesized that the P74L mutation impairs CLDN4-dependent TJ integrity, increasing paracellular calcium permeability and failing to prevent electrochemically driven calcium back-flux into the tubular lumen, thereby creating conditions permissive to hypercalciuric kidney stone disease. This study aimed to characterize the molecular and functional consequences of the P74L mutation using inducible epithelial cell models expressing human wild-type (WT) or mutant CLDN4, providing new insight into how CLDN4 variants compromise epithelial integrity and contribute to kidney-stone formation.

## MATERIALS AND METHODS

### Patient Data and Ethical Approval

Clinical and genetic data for CLDN4 protein variants (p.P74L and p.A195T) were obtained from the BKSR, coordinated by Dr. Daniel Fuster (University of Bern, Switzerland). The study cohort, phenotypic assessments, and genetic analysis pipeline have been previously described in detail (36). The BKSR complies with the Declaration of Helsinki and received approval from the Kantonale Ethikkommission Bern (Approval No. BE 95/06). Written informed consent for genetic analysis and anonymized data sharing was obtained from all participants. Only de-identified data were shared with our group; therefore, no additional ethics approval was required for the present study. The two individuals carrying the heterozygous missense substitution cCg/cTg in position 427 in the cDNA (rs782390551) resulting in the P74L substitution in CLDN4, are unrelated males, with 1 and 6 kidney stone events, respectively. The first individual was 56 years old at diagnosis, displayed osteopenia, obesity grade 3 and a family history of kidney stones. The second individual was 33 years old at diagnosis, was hypercalciuric, also had a family history of kidney stones and had been previously diagnosed with a membranoproliferative glomerulonephritis by kidney biopsy in 1984. No genetic variation in other genes known to cause kidney stones were identified. Stones from both individuals were predominantly composed of either calcium oxalate or calcium phosphate.

### Cell Culture and Generation of Inducible CLDN4 Cell Lines

M1 mouse cortical CD epithelial cells (ATCC CRL-2038) were cultured in DMEM/F-12 (Gibco, Thermo Fisher Scientific; Cat. no. 11330-032) supplemented with 10% fetal bovine serum (Gibco; Cat. no. A5256701), 1% antibiotic–antimycotic (Cytiva HyClone; Cat. no. SV30010), and 2.4 mM L-glutamine (Agilent; Cat. no. 103579) and maintained at 37 °C in a humidified atmosphere containing 5% CO₂. Doxycycline (DOX)-inducible M1 cell lines were generated using the Tet-On 3G lentiviral expression system (Takara Bio Inc.). pLVX-TRE3G constructs encoding hemagglutinin (HA)-tagged human WT CLDN4 or the P74L CLDN4 variant were custom generated by VectorBuilder (Chicago, IL, USA). Lentiviral particles were produced in Lenti-X 293T cells using standard packaging procedures (Clontech Laboratories). M1 cells were first transduced with pLVX-Tet3G and selected with G418 (1 mg/mL; Gibco; Cat. no. 11811031), followed by transduction with lentiviruses encoding WT or P74L CLDN4 and selection with puromycin (4 µg/mL; Gibco; Cat. no. A11138-03). The resulting stable cell lines permit doxycycline (DOX)-dependent expression of HA-tagged CLDN4 variants. Inducible expression was confirmed by immunoblotting and immunofluorescence following DOX treatment (1 µg/mL for 24 h; MilliporeSigma; Cat. no. D3072).

### Ussing Chamber Experiments

M1 cells expressing DOX-inducible WT or P74L CLDN4 were seeded on Snapwell permeable filter inserts (Corning; 12-mm diameter, 0.4-µm pore polycarbonate membrane; Cat. no. 3407) and cultured for approximately 10 days to allow formation of a polarized epithelial monolayer. Transepithelial electrical resistance (TER) was monitored periodically during culture using an epithelial voltohmmeter (MilliporeSigma). Once stable TER values indicative of polarization were achieved, cells were induced with DOX for 24 h as indicated and mounted in Ussing chambers (EasyMount diffusion chambers; Physiological Instruments, San Diego, CA, USA) enabling electrophysiological measurements.

Transepithelial potential difference (PD, mV) was recorded continuously using a voltage/current clamp (EVC4000; World Precision Instruments, Sarasota, FL, USA), and TER (Ω·cm²) was determined by applying brief ±90 µA current pulses and calculating resistance using Ohm’s law, with values normalized to membrane surface area. Signals were acquired using Chart software (version 4.2; ADInstruments).

To assess relative sodium and chloride permeability, a dilution potential was generated by replacing the apical solution with a low-sodium solution (30 mM Na⁺, full solution compositions in **Table S3**.) while maintaining the basolateral compartment in control solution. After stabilization, the apical solution was returned to control conditions. Paracellular calcium permeability was assessed using a calcium bi-ionic dilution protocol as previously described (41), in which the apical solution was replaced with a high-calcium solution (70 mM Ca²⁺) and the basolateral compartment was simultaneously exchanged with a phosphate-free control solution. Changes in PD were recorded after reaching a stable peak and again following restoration of control solutions.

Forskolin (10 µM) was added at the end of each experiment to confirm epithelial viability and responsiveness. Liquid junction potentials arising from solution substitutions were determined using empty filter inserts processed in parallel and were used to correct all measured PD values. Corrected PD and TER values were used to calculate absolute sodium and chloride permeabilities (P_Na_ and P_Cl_, 10^-4^ cm/s), their relative permeability ratios (P_Na_/P_Cl_) and calcium permeability parameters (P_Ca_, 10^-4^ cm/s and P_Ca_/P_Na_) using established equations (42). All solutions were adjusted to physiological pH (7.4) and osmolality (290–292 mOsm/kg).

#### 1. Statistical Analysis

Statistical analyses were performed using GraphPad Prism (version 10.4.1). Data are presented as mean ± SEM, with each symbol representing an independent biological replicate. Statistical tests, including unpaired two-tailed Student’s *t*-tests and one- or two-way ANOVA with Tukey’s multiple comparisons test, were applied as indicated in the figure legends. Normalization strategies and sample sizes for each experiment are described in the corresponding figure legends. Exact *p* values are reported where applicable; statistical significance was defined as *p* < 0.05.

**Additional detailed experimental procedures are provided in the *Supplementary Material*.**

## RESULTS

### CLDN4 variants are identified in patients with recurrent calcium nephrolithiasis

To identify novel genes contributing to idiopathic calcium nephrolithiasis, whole-exome sequencing and biochemical data analyzed from 787 kidney stone disease adults from the BKSR uncovered two un-related male patients who carried the heterozygous missense substitution cCg/cTg in position 427 in the *CLDN4* cDNA (rs782390551), resulting in the P74L substitution in CLDN4 protein (**Table 1**) (36). This variation has an allele frequency of 0.00005174 (gnomAD) and was assessed as “possibly damaging” (0.903) by PolyPhen and deleterious by SIFT. The P74L variant affects a conserved residue within the ECL1, a region critical for paracellular ion selectivity (21,37–40,43,44). Both patients reported a family history of KS and showed evidence of osteopenia.

**Table 1.** Genetic, biochemical, and clinical characteristics of two male patients from the Bern Kidney Stone Registry carrying missense variants in CLDN4. Amino acid substitutions and corresponding codon changes are shown in the mutation column. Plasma calcium concentrations are within the normal reference range (2.2–2.6 mmol/L) for both individuals, whereas urinary calcium excretion varied substantially among patients (normal range for adult males: <7.5 mmol/day). Both individuals reported a family history of KS and exhibited osteopenia.

| Patient | Sex | Stone events | CLDN4 mutation (protein, codon) | Pathogenicity | Plasma Ca<sup>2+</sup> (mmol/L) | Urinary Ca<sup>2+</sup> (mmol/day) | Family history / Other issues |
| --- | --- | --- | --- | --- | --- | --- | --- |
| 1 | Male | 1 | P74L (cCg → cTg) | Likely Pathogenic | 2.2 | 4.2 | Family history of KS; Osteopenia |
| 2 | Male | 6 | P74L (cCg → cTg) | Likely Pathogenic | 2.4 | 14.5 | Family history of KS; Osteopenia; Previous glomerular nephritis |

Plasma calcium concentrations were within the normal reference range in both individuals, but urinary calcium excretion varied substantially, ranging from normal (individual 1) to hypercalciuric (individual 2). Given this variant’s frequency, predicted pathogenicity, and location within the pore-forming region, we conducted mechanistic investigations on the P74L variant to assess its possible role in calcium homeostasis.

To locate the P74L substitution within the CLDN4 tri-dimensional structure, we mapped residue 74 onto the AlphaFold-predicted structural model of human CLDN4 (UniProt accession O14493) (**Figure 1**). In the WT protein (**Figure 1A**), Proline 74 lies within the ECL1. Visualization of the mutant (**Figure 1B**) shows the substitution of proline with leucine at the same position. Proline is conformationally restricted at the backbone due to its cyclic side chain, whereas standard aliphatic residues such as leucine exhibit greater conformational flexibility, possibly altering the protein conformation (45,46). Although no major rearrangement is apparent in this static model, the biochemical nature and structural location of the P74L substitution may affect dynamic conformational movement within a region relevant to CLDN4 function.

**Figure 1.**
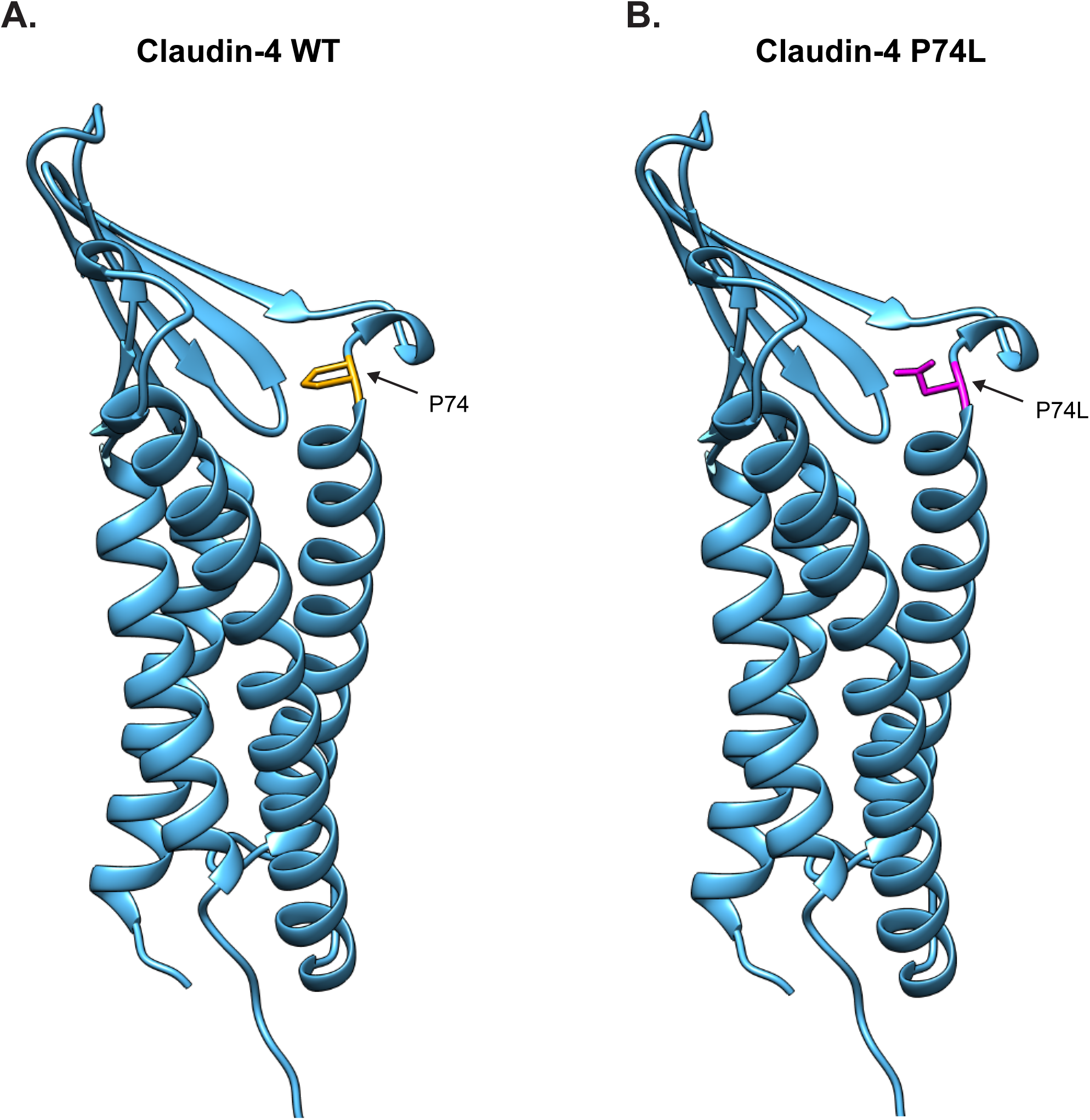
Structural visualization of wild-type (WT) and P74L Claudin-4 (CLDN4). (A) Structural model of WT CLDN4 highlighting proline 74 (P74, yellow) located within the first extracellular loop (ECL1). (B) Structural model of the P74L CLDN4 variant showing substitution of proline with leucine at position 74 (P74L, magenta). Residue 74 is shown in stick representation, with arrows indicating its location. Models were generated using UCSF Chimera.

### The P74L CLDN4 variant exhibits reduced steady-state abundance

To validate our inducible expression system, we examined hemagglutinin-tagged human claudin-4 (HA–hCLDN4) levels in WT and P74L CLDN4–expressing M1 cells under +DOX and –DOX conditions (**Figure 2A–B**). As expected, HA–hCLDN4 was undetectable in both cell lines in the absence of DOX. Following induction, WT cells exhibited robust HA–hCLDN4 expression, whereas P74L cells displayed significantly lower protein levels compared with WT under equivalent induction conditions (0.19 ± 0.01, n = 3, P74L +DOX, vs. 1.00 ± 0.05, n = 3, WT +DOX; *p* < 0.0001), indicating reduced steady-state abundance of the mutant protein.

**Figure 2.**
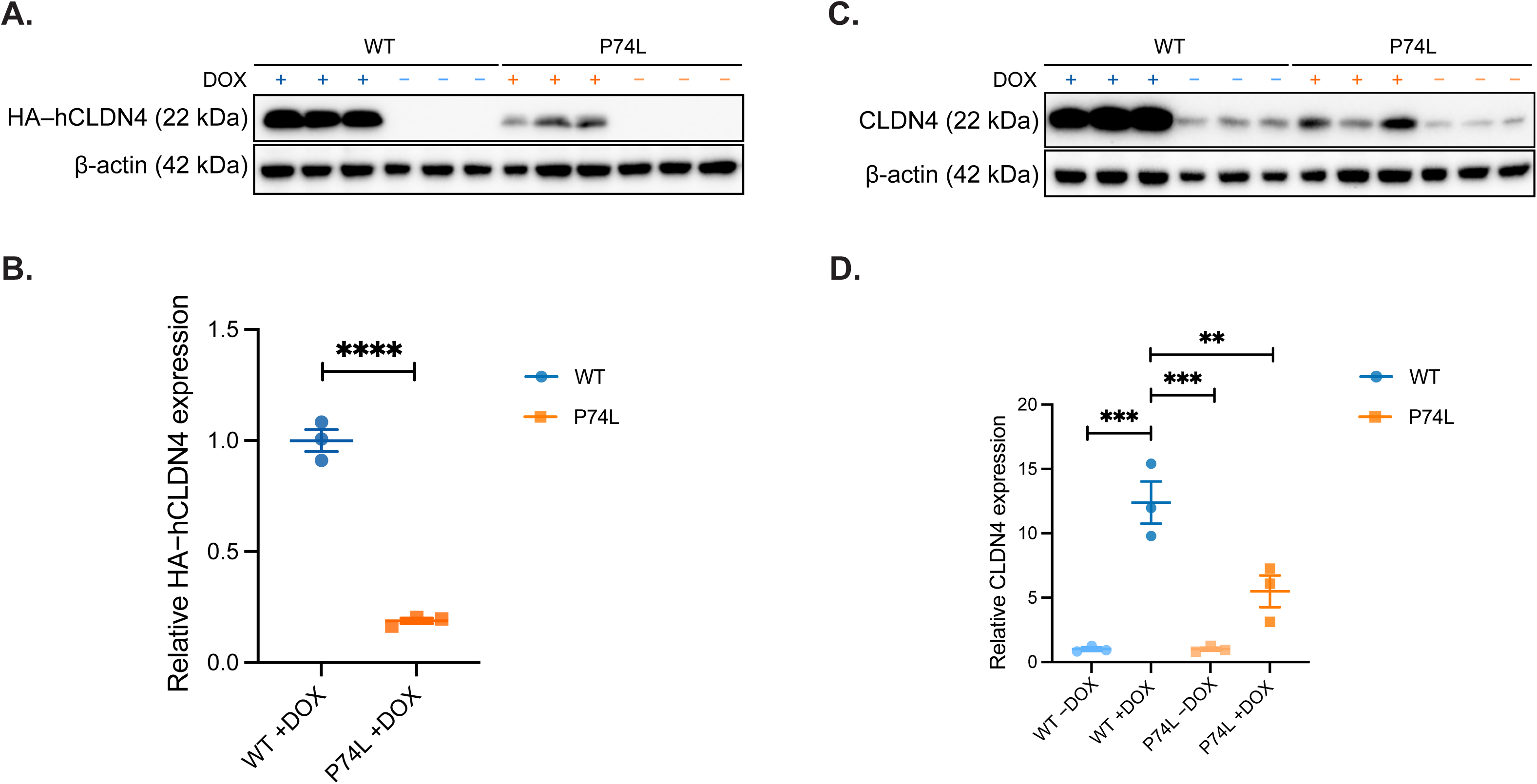
Immunoblot analysis and quantification of inducible human Claudin-4 (CLDN4) and total CLDN4 protein abundance in wild-type (WT) and P74L CLDN4–expressing M1 cells. (A) Immunoblots showing hemagglutinin-tagged human claudin-4 (HA–hCLDN4) expression in WT and P74L CLDN4–expressing M1 cells under +DOX and –DOX conditions (DOX, doxycycline). β-actin served as the loading control. (B) Quantification of HA–hCLDN4 protein levels normalized to β-actin and expressed relative to the average of the WT +DOX group. Data represent mean ± SEM from n = 3 independent experiments, with each symbol indicating a biological replicate. Statistical analysis was performed using an unpaired *t*-test (**** *p* < 0.0001). (C) Immunoblot of CLDN4 and β-actin in WT and P74L CLDN4–expressing M1 cells under +DOX and –DOX conditions. The anti-CLDN4 antibody used detects both murine and human CLDN4. (D) Quantification of CLDN4 normalized to β-actin and expressed relative to the −DOX average of each cell line. Data represent mean ± SEM from n = 3 independent experiments, with each symbol indicating a biological replicate. Statistical analysis was performed using two-way ANOVA with Tukey’s multiple comparisons test (** *p* = 0.0062; *** *p* = 0.0002).

To determine whether induction of human WT or P74L CLDN4 altered endogenous CLDN4 expression, we next assessed CLDN4 protein abundance using an anti-CLDN4 antibody that detects both the murine and human protein (**Figure 2C–D**). In WT cells, DOX induction was associated with a significant 12.39 ± 1.63 fold (mean ± SEM) increase in CLDN4. In contrast, P74L cells showed a non-significant 5.49 ± 1.23 fold (mean ± SEM) increase in CLDN4 with DOX treatment.

Together, these results demonstrate successful inducible expression of WT and P74L CLDN4, and reduced steady-state accumulation of the P74L variant in M1 cells.

### P74L CLDN4 displays reduced protein stability

To determine whether the reduced steady-state abundance of P74L CLDN4 reflects decreased protein stability, we performed cycloheximide (CHX) chase assays in DOX-induced WT and P74L CLDN4–expressing M1 cells (**Figure 3**). Following CHX treatment, HA–hCLDN4 levels declined progressively over time in both cell lines, with a markedly faster decay observed for P74L compared with WT CLDN4 (**Figure 3A–B**). Quantitative analysis (**Figure 3C**) revealed that P74L CLDN4 exhibited a markedly shorter half-life (t_₁⁄₂_ = 0.96 h) compared with WT CLDN4 (t_₁⁄₂_ = 4.74 h). These results indicate that the P74L variant exhibits reduced protein stability compared with WT CLDN4.

**Figure 3.**
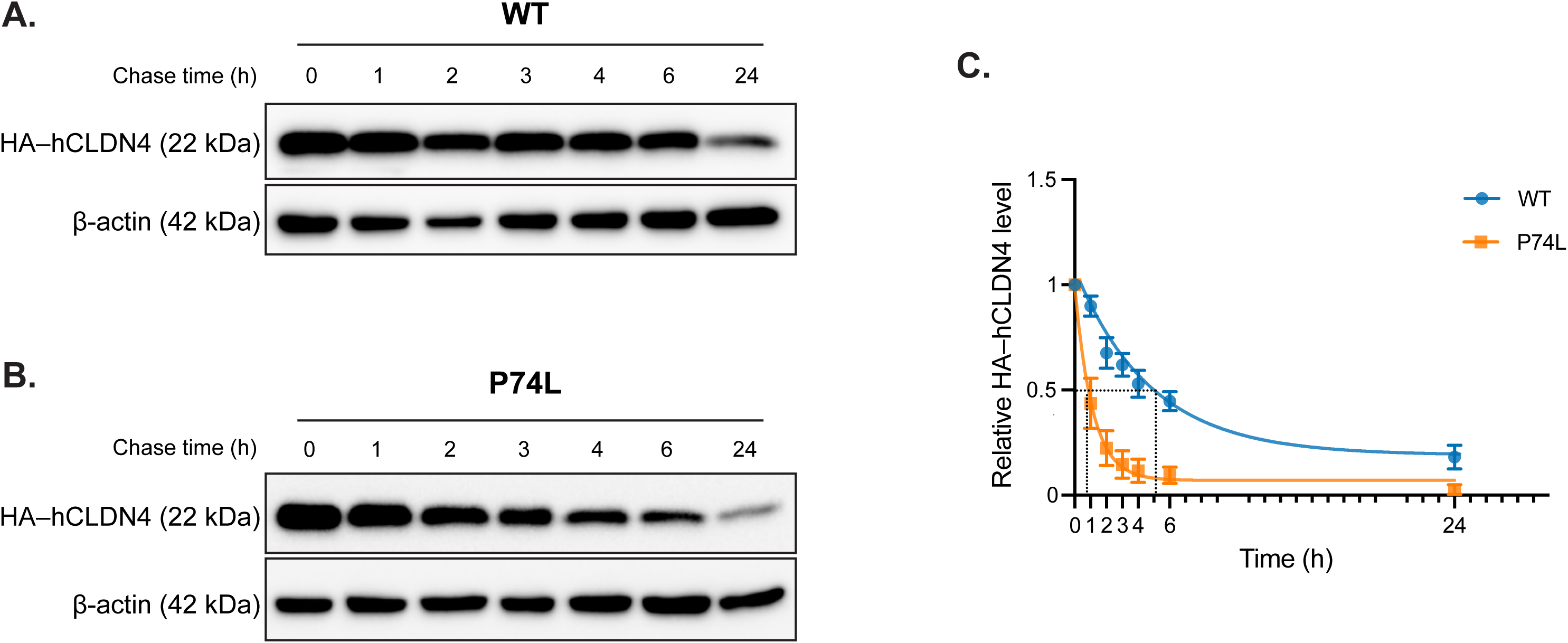
Cycloheximide (CHX) chase assay of wild-type (WT) and P74L Claudin-4 (CLDN4). Representative immunoblots of WT (A) and P74L (B) CLDN4–expressing M1 cells induced with doxycycline (DOX) and treated with CHX for 24h, then lysed at 0, 1, 2, 3, 4, 6, and 24 h post-CHX. Blots were probed for hemagglutinin-tagged human claudin-4 (HA–hCLDN4) and β-actin. (C) Quantification of HA–hCLDN4 protein levels during the CHX chase. Band intensities were normalized to β-actin and expressed relative to the corresponding 0 h value for each biological replicate (n = 3). Data are shown as mean ± SEM with one-phase exponential decay fits. A horizontal dotted line at *Y* = 0.5 denotes 50% of the initial protein level. The half-life (t₁⁄₂) for each condition was defined as the intersection of this line with the fitted decay curve; vertical dotted lines indicate these corresponding *X*-values for WT and P74L. Blots in (A, B) show one representative experiment, and the quantification in (C) summarizes three independent biological replicates (n = 3).

### P74L CLDN4 shows defective membrane localization and reduced surface expression

To examine whether reduced stability of P74L CLDN4 is associated with altered subcellular localization and surface expression, we performed cell-surface biotinylation assays in DOX-induced cells (**Figure 4A**). WT CLDN4 was readily detected in the biotinylated fraction, whereas P74L CLDN4 exhibited significantly reduced surface abundance. Quantification of the biotinylated fraction normalized to input confirmed a significant decrease in surface-expressed P74L CLDN4 to 45 ± 16 % of WT cell surface abundance (*p* = 0.0266; **Figure 4B**). To further assess protein localization, we analyzed the distribution and surface expression of WT and P74L CLDN4 in polarized M1 cells. Immunofluorescence revealed that WT CLDN4 localized predominantly to cell–cell contacts and colocalized with the TJ marker ZO-1 following DOX induction (**Figure 4C**). In contrast, P74L CLDN4 exhibited reduced junctional localization and a more diffuse intracellular staining pattern under the same conditions (**Figure 4C**). No HA–hCLDN4 signal was detected in either cell line under −DOX conditions.

**Figure 4.**
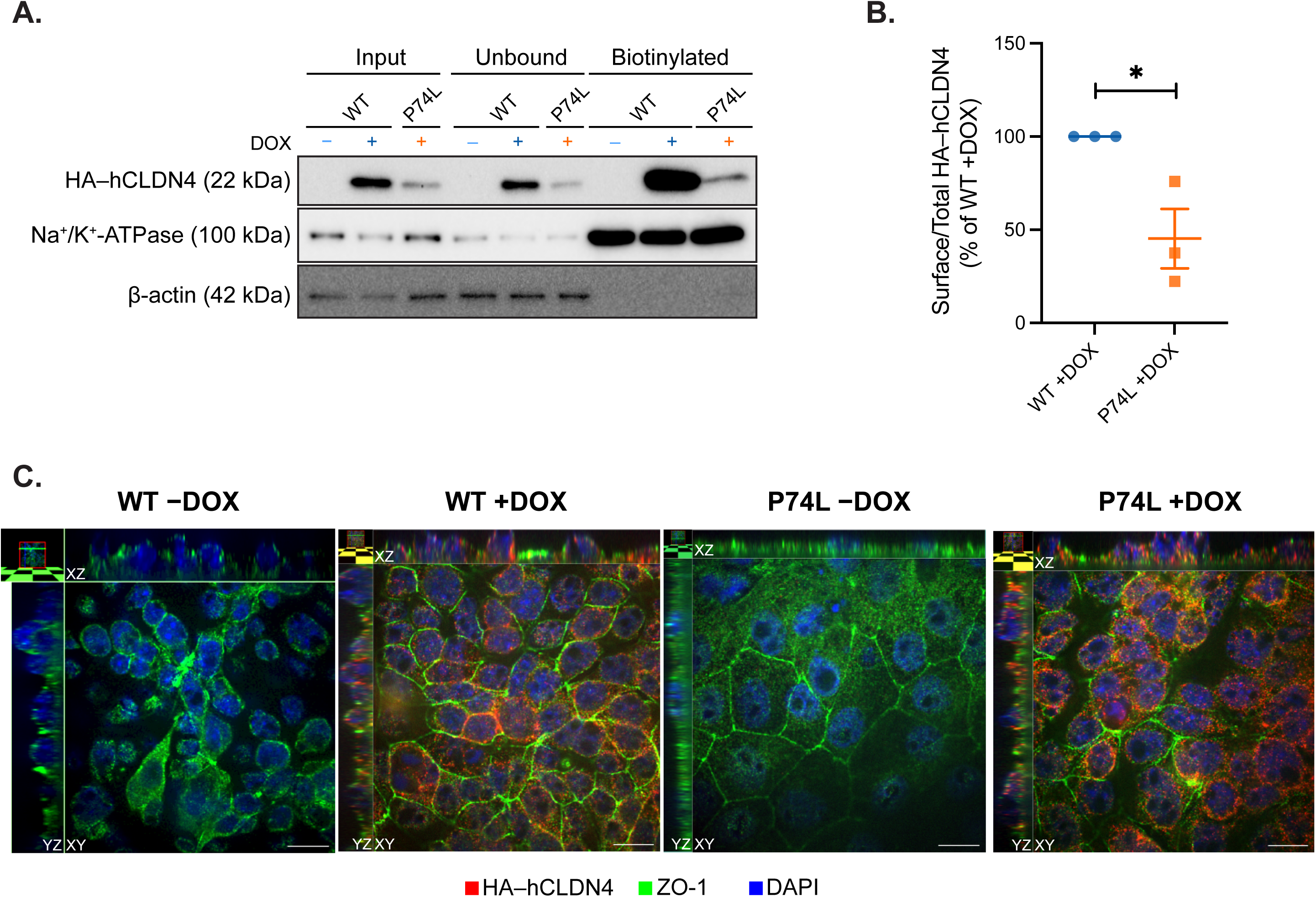
Surface expression and localization of wild-type (WT) and P74L Claudin-4 (CLDN4). (A) Representative immunoblot of cell-surface biotinylation showing Input, Unbound, and Biotinylated fractions from WT –DOX, WT +DOX, and P74L +DOX M1 cells. Blots were probed for HA–hCLDN4, Na⁺/K⁺-ATPase (surface control), and β-actin (cytosolic control). (B) Quantification of surface HA–hCLDN4. Band intensities in the Biotinylated fraction were normalized to the corresponding Input for each biological replicate and expressed relative to the WT +DOX condition within the same replicate. Data are presented as mean ± SEM from n = 3 independent experiments, with each symbol representing one biological replicate. Statistical significance was determined using an unpaired *t*-test (* *p* = 0.0266). (C) Immunofluorescence of polarized WT and P74L CLDN4–expressing M1 cells grown on Transwell filters under –DOX and +DOX conditions (DOX). Cells were stained for hemagglutinin-tagged human claudin-4 (HA–hCLDN4; red), ZO-1 (green), and DAPI (blue). Images are Z-stack acquisitions, with the central XY panel accompanied by orthogonal XZ (top) and YZ (left) views. Scale bars correspond to 10 μm.

Together, these data demonstrate that the P74L variant exhibits impaired membrane localization and reduced surface expression relative to WT CLDN4.

### WT and P74L CLDN4 expression differentially alter TJ CLDN composition

To determine whether inducible expression of WT or P74L CLDN4 alters TJ CLDN composition through transcriptional or post-transcriptional mechanisms, we first assessed endogenous mouse *Cldn4* and *Cldn8* mRNA expression by quantitative reverse transcription polymerase chain reaction (qRT-PCR). No significant differences were detected across cell lines or DOX conditions (**Figure 5A&B**), indicating that inducible CLDN4 expression does not elicit detectable transcriptional changes in these genes.

**Figure 5.**
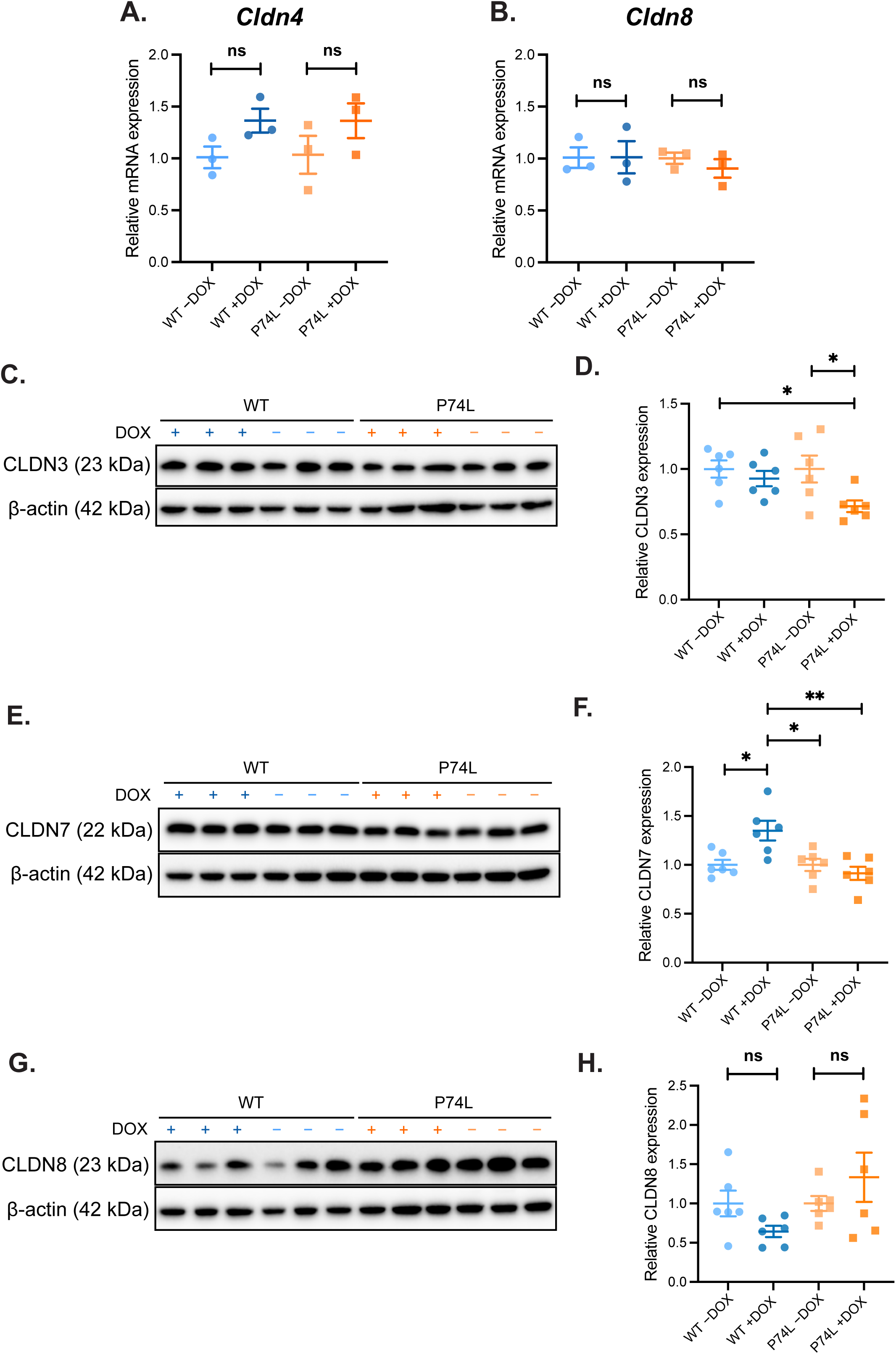
Expression of claudin (CLDN) proteins in wild-type (WT) and P74L Claudin-4 (CLDN4)–expressing M1 cells under +DOX and −DOX conditions. Relative mRNA expression of endogenous mouse *Cldn4* (A) and *Cldn8* (B) in WT and P74L CLDN4–expressing M1 cells under −DOX and +DOX conditions. Expression levels were normalized to *Rplp0* and expressed relative to the corresponding −DOX baseline for each cell line using the 2^−ΔΔCT^ method. Data represent mean ± SEM from n = 3 independent experiments, with each symbol indicating a biological replicate. Statistical comparisons were performed using two-way ANOVA with Tukey’s multiple comparisons test; no significant cell line– or DOX-dependent differences were detected (ns, not significant). (C, E, G) Representative immunoblots for CLDN3, CLDN7, and CLDN8, respectively, with β-actin as a loading control, each lane corresponding to a biological replicate. Blots shown are from one of two independent experiments (n = 3 per blot; total n = 6 biological replicates per condition). (D, F, H) Quantification of CLDN3, CLDN7, and CLDN8 protein levels normalized to β-actin and expressed relative to the −DOX average baseline for each cell line. Data represent mean ± SEM from n = 6 biological replicates. Statistical analysis for CLDN3 and CLDN7 was performed using two-way ANOVA with Tukey’s multiple comparisons test. For CLDN3, significant differences were observed between WT −DOX and P74L +DOX (* *p* = 0.0480) and between P74L −DOX and P74L +DOX (* *p* = 0.0480). For CLDN7, significant differences were detected between WT +DOX and WT −DOX (* *p* = 0.0142), between WT +DOX and P74L −DOX (* *p* = 0.0142), and between WT +DOX and P74L +DOX (** *p* = 0.0021). For CLDN8, neither two-way nor one-way ANOVA detected significant differences between conditions or cell lines. ns, not significant.

We next examined whether inducible WT or P74L CLDN4 expression alters the abundance of other TJ CLDNs at the protein level. Immunoblot analysis revealed detectable expression of CLDN3, CLDN7, and CLDN8 across conditions, with distinct patterns depending on the claudin examined (**Figure 5C, E, G**). Analysis of band intensity showed that CLDN3 protein levels were not significantly altered by CLDN4 induction in WT cells (**Figure 5D**); however, CLDN3 abundance was significantly reduced following induction in P74L CLDN4–expressing cells (0.72 ± 0.04, n = 6, P74L +DOX, vs. 1.00 ± 0.10, n = 6, P74L −DOX ; *p* = 0.0480) and was also significantly lower than WT cells under −DOX conditions (0.72 ± 0.04, n = 6, P74L +DOX, vs. 1.00 ± 0.07, n = 6, WT −DOX; *p* = 0.0480). CLDN4 induction resulted in a significant increase in CLDN7 protein abundance in WT cells (1.35 ± 0.10, n = 6, WT +DOX, vs. 1.00 ± 0.05, n = 6, WT−DOX; *p* = 0.0142), whereas no induction-dependent change was observed in P74L cells (**Figure 5F**). Moreover, while CLDN7 abundance significantly increased in +DOX WT cells compared to -DOX WT cells, a similar increase was not observed in +DOX P74L versus -DOX P74L cells. Finally, CLDN8 protein levels were not significantly affected by CLDN4 induction in either WT–or P74L–expressing cells despite a trend for increase in +DOX P74L mutant cells versus P74L – DOX cells, and no significant differences were detected between cell lines (**Figure 5H**).

Given that the CLDN4 variants were identified in patients with calcium-based kidney stones, we further assessed whether inducible CLDN4 expression is associated with broader transcriptional changes in distal epithelial calcium transport and homeostasis pathways. We examined the mRNA expression of selected calcium transport–related genes *Trpv5*, *Trpv6*, *Calb1*, *Atp2b1*, and *Slc8a1* (2,31–34,47,48). No significant cell line– or DOX-dependent differences were detected for any of the targets examined (**Supplementary Figure 1)**. In addition, because paracellular chloride permeability mediated by CLDN4 functionally interfaces with electrogenic sodium and potassium transport in the distal nephron, we also assessed the expression of the genes encoding ENaC (*Scnn1g*) and the renal outer medullary potassium channel (ROMK; *Kcnj1*) (21,49,50). Again, no significant cell line– or DOX-dependent differences were detected (**Supplementary Figure 2**).

Together, these data indicate that inducible CLDN4 expression does not produce detectable transcriptional changes in endogenous Cldn4/Cldn8 or distal ion transport genes in M1 cells, while TJ composition differs selectively at the protein level between WT and P74L CLDN4–expressing cells.

### P74L CLDN4 fails to increase transepithelial electrical resistance (TER) and to restrict ion permeability

To obtain a functional readout of epithelial barrier properties and TJ–dependent paracellular conductance associated with the P74L variant, we measured the transepithelial electrical resistance (TER) and ion permeabilities in WT versus P74L CLDN4–expressing polarized M1 monolayers (51,52).

Expression of WT CLDN4 resulted in a significant increase in TER compared with WT −DOX monolayers (2.70 ± 0.23, n = 8, WT +DOX, vs. 1.00 ± 0.13, n = 6, WT −DOX; *p* < 0.0001; **Figure 6A**). In contrast, induction of P74L CLDN4 did not significantly change TER, despite a slight trend for increase. Given the established role of CLDN4 in paracellular chloride permeability and its functional coupling to sodium transport, we next examined sodium and chloride permeability parameters (21,29). The permeability ratio of sodium to chloride (P_Na_/P_Cl_) was not significantly altered by expression of either WT or P74L CLDN4 (**Figure 6B**). However, induction of WT CLDN4 significantly reduced sodium permeability (P_Na_) relative to WT −DOX monolayers (0.36 ± 0.03, n = 8, WT +DOX, vs. 1.00 ± 0.14, n = 6, WT −DOX; *p* < 0.0001; **Figure 6C**), whereas P74L CLDN4 expression failed to reduce P_Na_. Accordingly, P_Na_ values in both P74L −DOX and P74L +DOX monolayers were significantly higher than those observed in WT +DOX monolayers (1.00 ± 0.06, n = 6, P74L −DOX, vs. 0.36 ± 0.03, n = 8, WT +DOX *; p* < 0.0001; and 0.73 ± 0.8, n = 8, P74L +DOX, vs. 0.36 ± 0.03, n = 8, WT +DOX*; p* = 0.0100, respectively). Similarly, chloride permeability (P_Cl_) was significantly reduced upon WT CLDN4 expression (0.36 ± 0.03, n = 8, WT +DOX, vs. 1.00 ± 0.14, n = 6, WT −DOX; *p* = 0.0001) but was not altered by induction of P74L CLDN4 (**Figure 6D**). Both −DOX and +DOX P74L monolayers exhibited significantly higher P_Cl_ compared with WT +DOX monolayers (1.00 ± 0.07, n = 6, P74L −DOX, vs. 0.36 ± 0.03, n = 8, WT +DOX*; p* = 0.000; and 0.71 ± 0.8, n = 8, P74L +DOX, vs. 0.36 ± 0.03, n = 8, WT +DOX*; p* = 0.0201, respectively).

**Figure 6.**
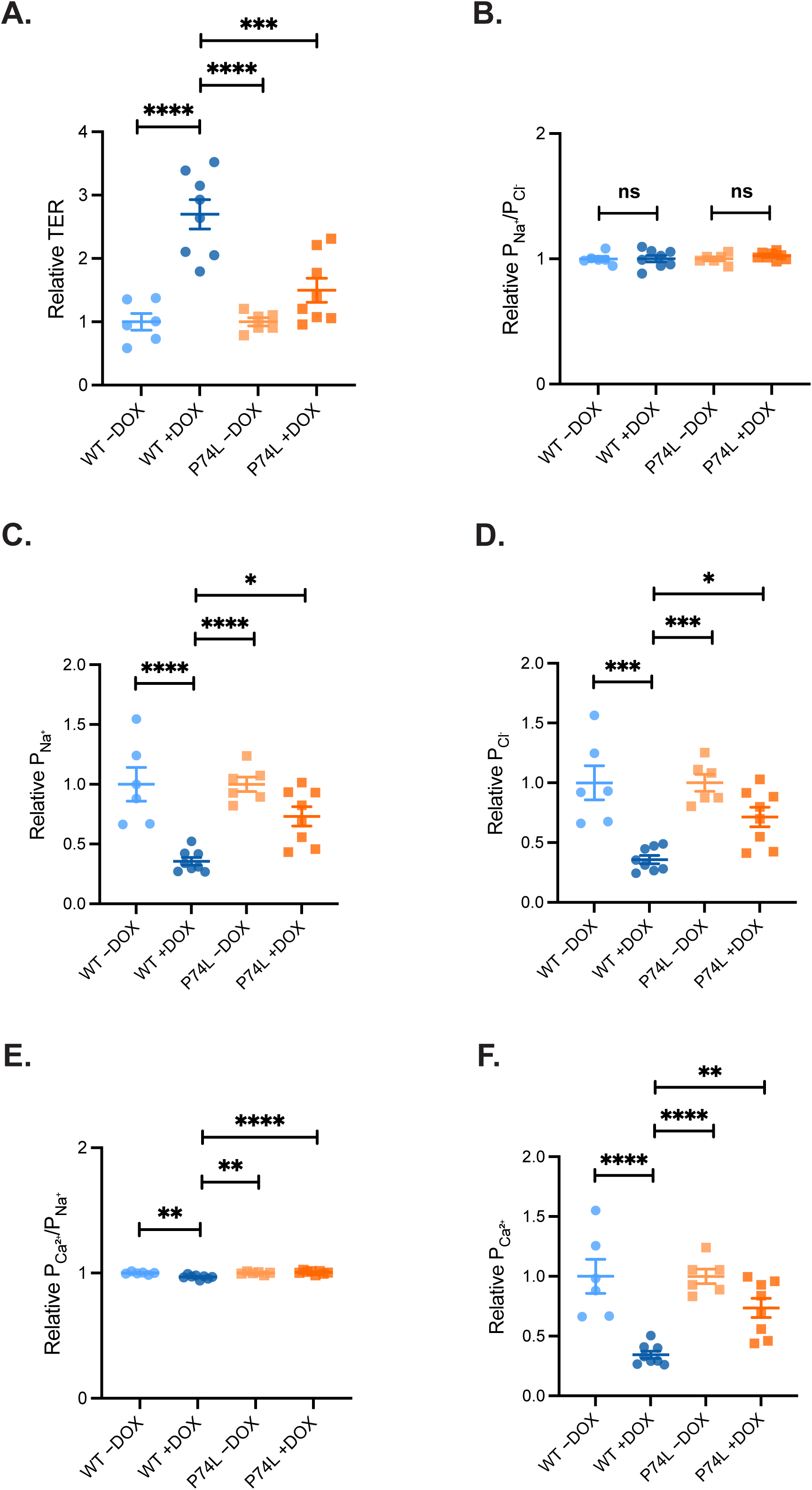
Transepithelial electrical resistance (TER) and ion permeabilities in wild-type (WT) and P74L Claudin-4 (CLDN4)–expressing M1 monolayers. (A) Relative TER and ion permeabilities measured in CLDN4 WT and P74L CLDN4–expressing M1 cell monolayers under −DOX and +DOX conditions: (B) P_Na_^+^/P_Cl_^-^; (C) P_Na_^+^; (D) P_Cl_^-^; (E) P_Ca_^2+^/P_Na_^+^; and (F) P_Ca_^2+^. P_Na_^+^, sodium absolute permeability; P_Cl_^-^, chloride absolute permeability; P_Ca_^2+^, calcium absolute permeability; P_Na_^+^/P_Cl_^-^ and P_Ca_^2+^/P_Na_^+^ indicate permeability ratios. Each point represents an independent experiment (−DOX: n = 6; +DOX: n = 8 per cell line). All values were normalized to the corresponding −DOX average baseline for each cell line. Data represent mean ± SEM, and statistical comparisons were performed using two-way ANOVA with Tukey’s multiple comparisons test. For TER (A), significant differences were observed between WT +DOX and WT −DOX (**** *p* < 0.0001) and between WT +DOX and P74L −DOX (**** *p* < 0.0001), as well as between WT +DOX and P74L +DOX (*** *p* = 0.0003). For P_Na_^+^ (C), significant differences were detected between WT +DOX and WT −DOX (**** *p* < 0.0001) and between WT +DOX and P74L −DOX (**** *p* < 0.0001), and between WT +DOX and P74L +DOX (* *p* = 0.0100). For P_Cl_^-^ (D), significant differences were observed between WT +DOX and WT −DOX (*** *p* = 0.0001) and between WT +DOX and P74L −DOX (*** *p* = 0.0001), as well as between WT +DOX and P74L +DOX (* *p* = 0.0201). For the P_Ca_^2+^/P_Na_^+^ (E), significant differences were detected between WT +DOX and WT −DOX (** *p* = 0.0011) and between WT +DOX and P74L −DOX (** *p* = 0.0011), and between WT +DOX and P74L +DOX (**** *p* < 0.0001). For P_Ca_^2+^ (F), significant differences were observed between WT +DOX and WT −DOX (**** *p* < 0.0001) and between WT +DOX and P74L −DOX (\*\*\*\**p* < 0.0001), as well as between WT +DOX and P74L +DOX (** *p* = 0.0072). ns, not significant.

Finally, because CLDN4 variants were identified in patients with calcium nephrolithiasis, we assessed calcium permeability to determine whether WT and P74L CLDN4 expression differentially affect paracellular calcium transport in the M1 cell line. Expression of WT CLDN4 significantly reduced the calcium-to-sodium permeability ratio (P_Ca_/P_Na_) compared with WT−DOX monolayers (0.97 ± 0.00, n = 8, WT +DOX, vs. 1.00 ± 0.00, n = 6, WT −DOX; *p* = 0.0010; **Figure 6E**). Consistent with this effect, absolute calcium permeability (P_Ca_) was also significantly reduced following WT CLDN4 expression (0.34 ± 0.03, n = 8, WT +DOX, vs. 1.00 ± 0.14, n = 6, WT −DOX; *p* < 0.0001; **Figure 6F**). In contrast, P74L CLDN4 expression neither significantly reduced P_Ca_/P_Na_ nor P_Ca_.

Together, these data indicate that WT CLDN4 expression enhances epithelial barrier properties and restricts paracellular ion permeability, whereas the P74L variant fails to confer these functional effects in M1 cells.

## DISCUSSION

In this study, we investigated the molecular and functional consequences of a kidney stone–associated CLDN4 variant, P74L, using inducible epithelial cell models expressing human WT or mutant CLDN4. Our findings demonstrate that the P74L substitution compromises CLDN4 protein stability, plasma membrane abundance, and TJ–dependent barrier function, resulting in a failure to restrict ion permeability. Collectively, these data identify P74L as a loss-of-function CLDN4 variant and provide mechanistic insight into how disruption of CLDN4-dependent paracellular regulation may loosen the distal epithelial barrier, potentially leading to calcium nephrolithiasis.

The focus on the P74L variant was motivated by its identification in two unrelated individuals with nephrolithiasis and a family history of kidney stones from the Bern Kidney Stone Registry. Additionally, this variant is localized within the ECL1 of CLDN4 (37–40), a conserved domain that governs paracellular ion selectivity among CLDNs and whose alterations modify epithelial barrier properties (53,54).

Using an inducible expression system, we observed that the P74L CLDN4 accumulated at significantly lower steady-state levels than WT CLDN4 despite equivalent transcriptional induction, indicating post-transcriptional regulation of protein abundance. This reduction was accompanied by a marked decrease in protein stability, as demonstrated by cycloheximide (CHX) chase analysis, consistent with an accelerated turnover of the mutant protein. Similar post-translational destabilization has been reported for disease-associated missense mutations in other nephron-expressed CLDNs, including CLDN16 variants linked to hereditary hypercalciuria (55). Together, these findings suggest that reduced protein stability is a primary mechanism underlying the diminished abundance of P74L CLDN4.

In addition to reduced protein stability, the P74L variant exhibited a pronounced defect in membrane localization and incorporation into the TJ complex. Whereas WT CLDN4 was readily detected at the cell surface and partially colocalized with ZO-1, P74L CLDN4 showed significantly reduced surface abundance and decreased junctional enrichment, indicating impaired functional availability at the epithelial barrier. Proper TJ localization is a critical determinant of CLDN function, as CLDNs must assemble to TJ strands to establish effective epithelial barrier properties (54,56). Consistently, disease-associated missense mutations in other nephron-expressed CLDNs, including CLDN16 and CLDN19, disrupt protein trafficking or assembly and impair paracellular transport function (55,57,58). Thus, the combined defects in stability and localization provide a mechanistic basis for the impaired barrier function observed with P74L CLDN4.

Consistent with the established role of CLDN4 in restricting paracellular sodium permeability and increasing epithelial resistance (21,59,60), inducible expression of WT CLDN4 markedly increased TER and reduced absolute paracellular permeability to sodium, chloride, and calcium, whereas the P74L variant failed to confer these effects. The effect of WT CLDN4 on chloride was unexpected as CLDN4 is permeable to chloride (61). Increased calcium permeability upon WT CLDN4 expression is also unlikely to be conducted by CLDN4 as it is not permeable to divalent ions (61). We thus wondered whether WT CLDN4 expression affected the expression of other CLDN proteins or transporters. In our study, expression of WT CLDN4 was associated with increased cation and anion pore CLDN7 abundance, with no change in general barrier CLDN3 or chloride pore, cation blocker CLDN8 abundance. An increase in CLDN7 does not explain the restricted permeability to chloride. Therefore, we speculate that other un-tested CLDN may restrict ion permeability in these cells or that protein abundance may not directly reflect correct localization/function. In contrast, P74L CLDN4 inducible expression led to a decrease in general barrier CLDN3 abundance with no other CLDN change, but failed to induce significant change in epithelial resistance and permeability when compared to -DOX P74L cells. This finding again reflects that protein abundance of selected CLDN does not fully capture the TJ status of these cells. Importantly, these protein-level differences occurred in the absence of detectable changes in endogenous *Cldn4* or *Cldn8* mRNA expression, or in genes involved in transcellular calcium handling and electrogenic sodium and potassium transport, indicating that inducible CLDN4 expression does not engage transcriptional regulation of major distal epithelial ion transport pathways. Instead, these findings point to post-transcriptional or junctional mechanisms governing selective TJ remodeling.

Finally, given that the two individuals carrying the P74L variation exhibited nephrolithiasis, our study aimed to examine the role of this variant in calcium transport. In the distal nephron, calcium reabsorption occurs via a transcellular pathway involving TRPV5, Calbindin D28K, basolateral PMCAs and NCX1 (31–34). These processes take place in a nephron segment that exhibits a lumen-negative transepithelial voltage driven by electrogenic sodium reabsorption (35). Under such conditions, divalent cations such as Ca²⁺ are submitted to a strong electrochemical driving force favoring movement into the tubular lumen. In this context, CLDN4 functions as a cation-restrictive TJ component that limits paracellular Ca²⁺ permeability, thereby preventing voltage-driven paracellular Ca²⁺ backflux. Accordingly, impaired CLDN4-dependent barrier function, as observed with the P74L variant, would be predicted to permit electrochemically driven paracellular Ca²⁺ backflux into the lumen, increasing urinary calcium excretion. This mechanism provides a physiologically coherent explanation linking increased paracellular Ca²⁺ permeability to voltage-driven Ca²⁺ backflux and hypercalciuria observed in one patient reported here and in *Cldn4* knockout mice (62–64). Even modest increases in distal paracellular Ca²⁺ backflux may, over time, contribute to elevated urinary calcium excretion and negative calcium balance, potentially predisposing affected individuals to nephrolithiasis and skeletal demineralization.

In summary, our study delineates a mechanistic pathway by which the CLDN4 P74L variant compromises epithelial TJ function through reduced protein stability and impaired junctional incorporation, leading to selective remodeling of TJ components and failure to restrict paracellular ion flux. Although our experiments were conducted in cell models, they reveal fundamental properties of CLDN4 that are likely relevant to distal nephron physiology and the pathogenesis of calcium nephrolithiasis. Future studies using *in vivo* models and patient-derived tissues will be required to establish the integrative physiological consequences of CLDN4 dysfunction in the intact nephron. Collectively, these findings advance our understanding of paracellular barrier regulation in the kidney and provide a foundation for exploring TJ modulation in the context of calcium homeostasis and kidney stone disease.

## ACKNOWLEDGEMENTS

The authors thank Debbie D O’Neill for assistance with help in Ussing chamber experiments and gratefully acknowledge Kiera Smith from the Cell Imaging Facility at the University of Alberta for valuable technical support. The University of Alberta Advanced Cell Exploration Core Facility (RRID:SCR_019182) receives financial support from the Faculty of Medicine & Dentistry, the University Hospital Foundation, Striving for Pandemic Preparedness – The Alberta Research Consortium, and the Canada Foundation for Innovation.

F.C. was supported by a University of Alberta Graduate Recruitment Scholarship, a Graduate Student Award and the Kidney Foundation of Canada (25KHRG-1422220). G.E. received Graduate Student Engagement Scholarships, the Faculty of Medicine and Dentistry Delnor Scholarship, the Faculty of Medicine and Dentistry 75th Anniversary Award from the University of Alberta, and an Alberta Graduate Excellence Scholarship. This study was supported by funding from the Natural Sciences and Engineering Research Council of Canada (Discovery Grant RGPIN-2024-04972), the Canadian Institutes of Health Research (PJT-168871), and the Kidney Foundation of Canada (25KHRG-1422220) to E.C. D.G.F. was supported by the Swiss National Science Foundation (grants #33IC30_166785, 33IC30_229750, 320030-231405 and 320030-227578) and the Swiss National Center of Competence in Research (NCCR Kidney.CH).

---

**Figure S1.**
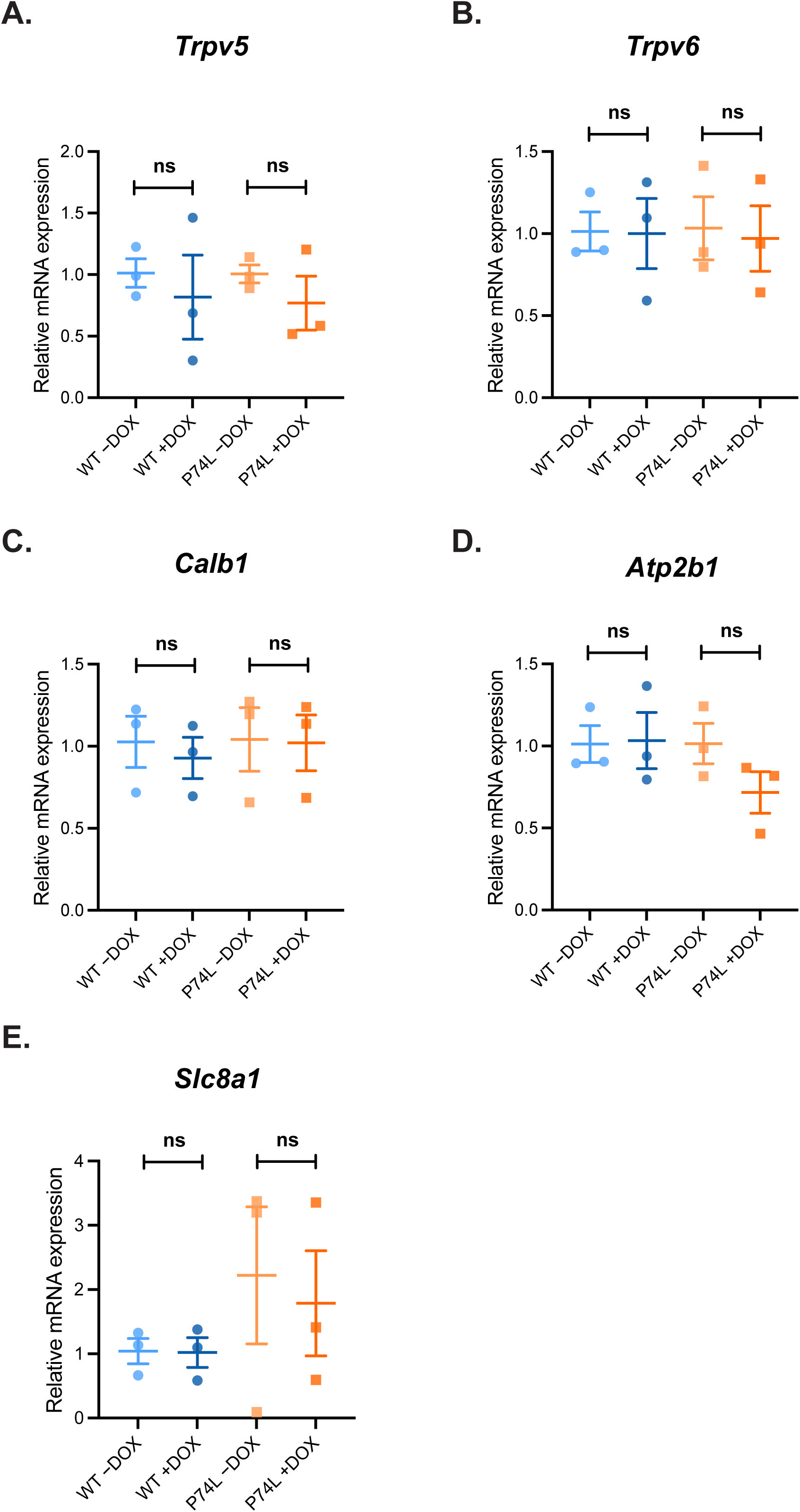

**Figure S2.**
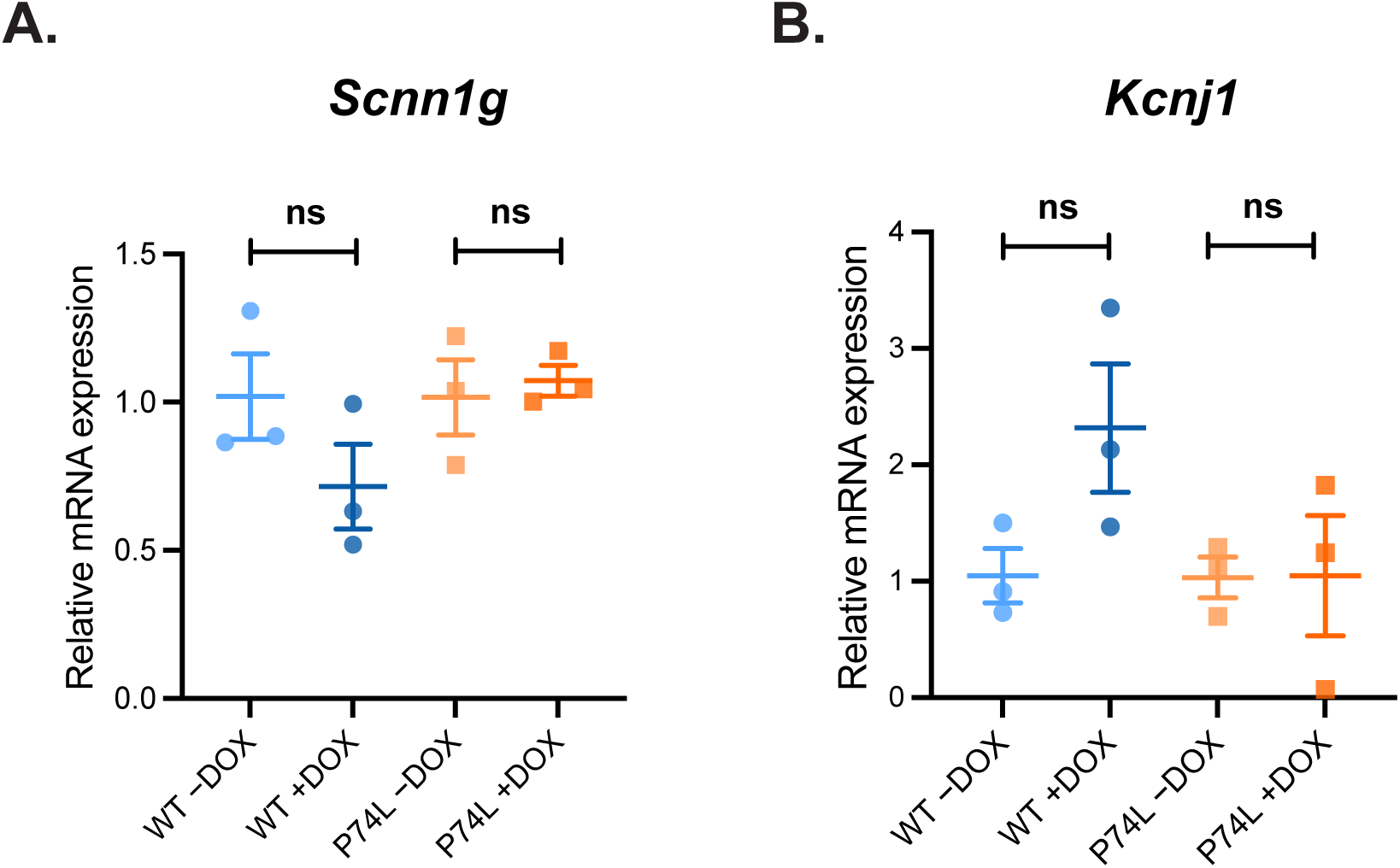

**Figure S3.**
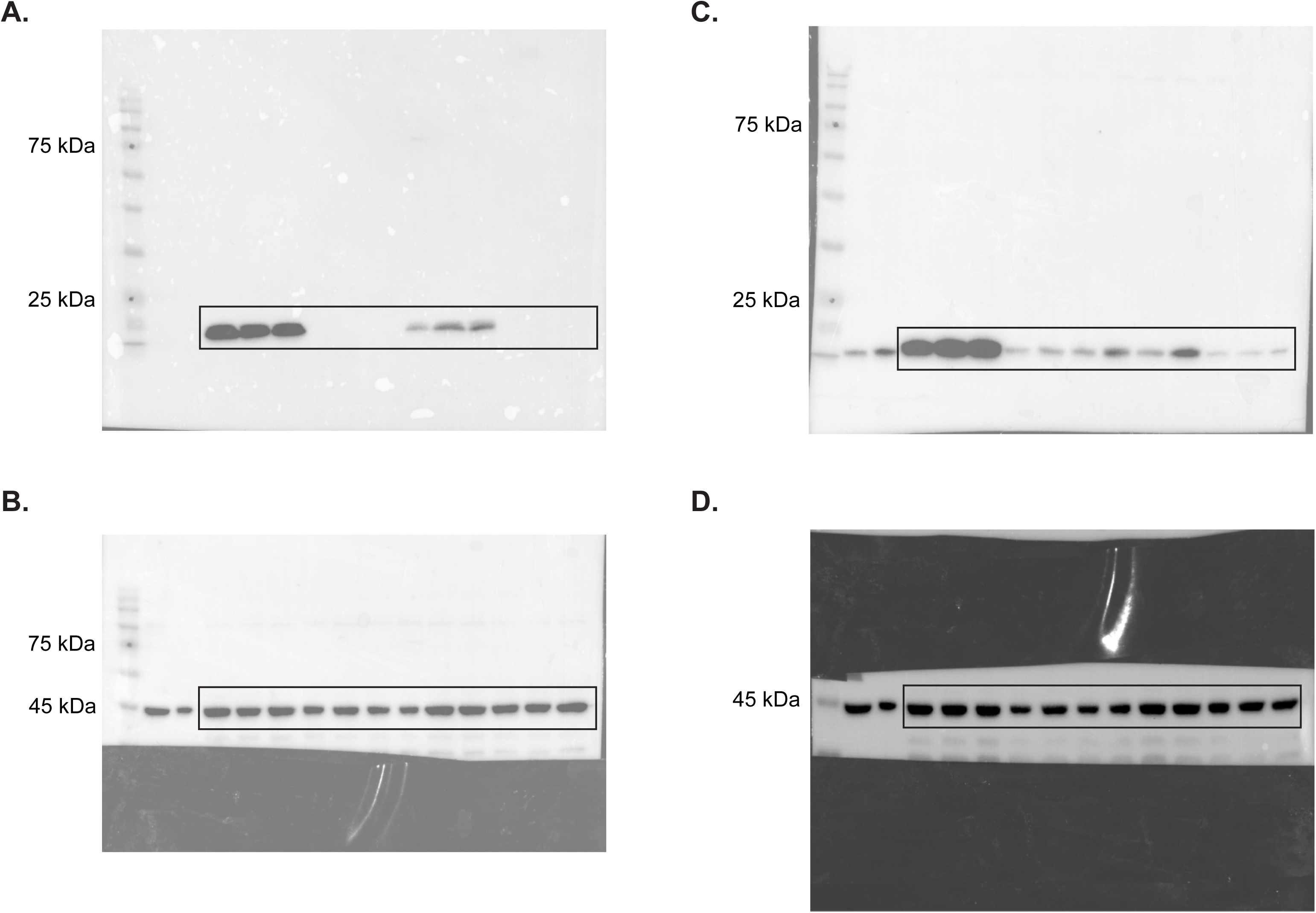

**Figure S4.**
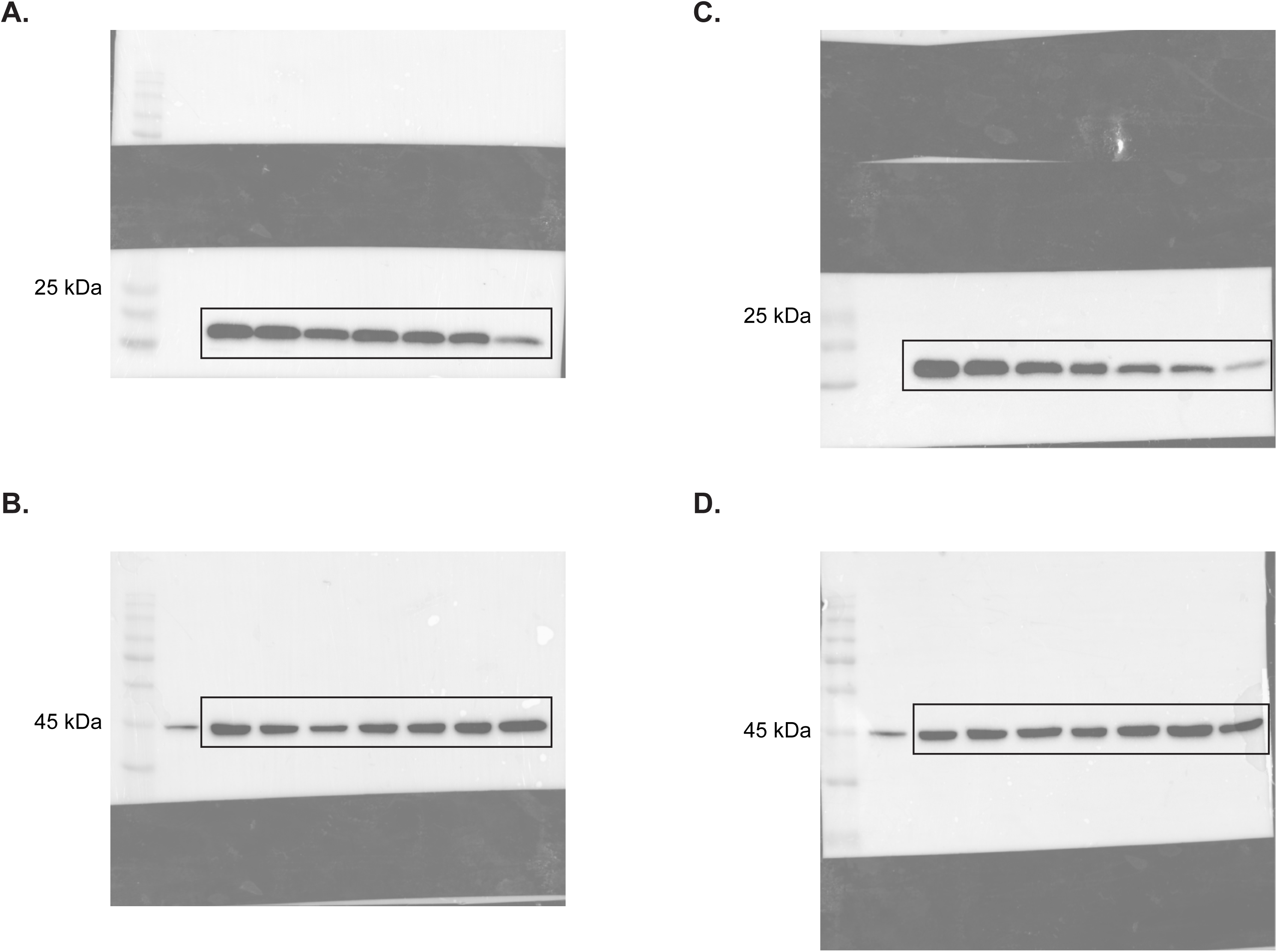

**Figure S5.**
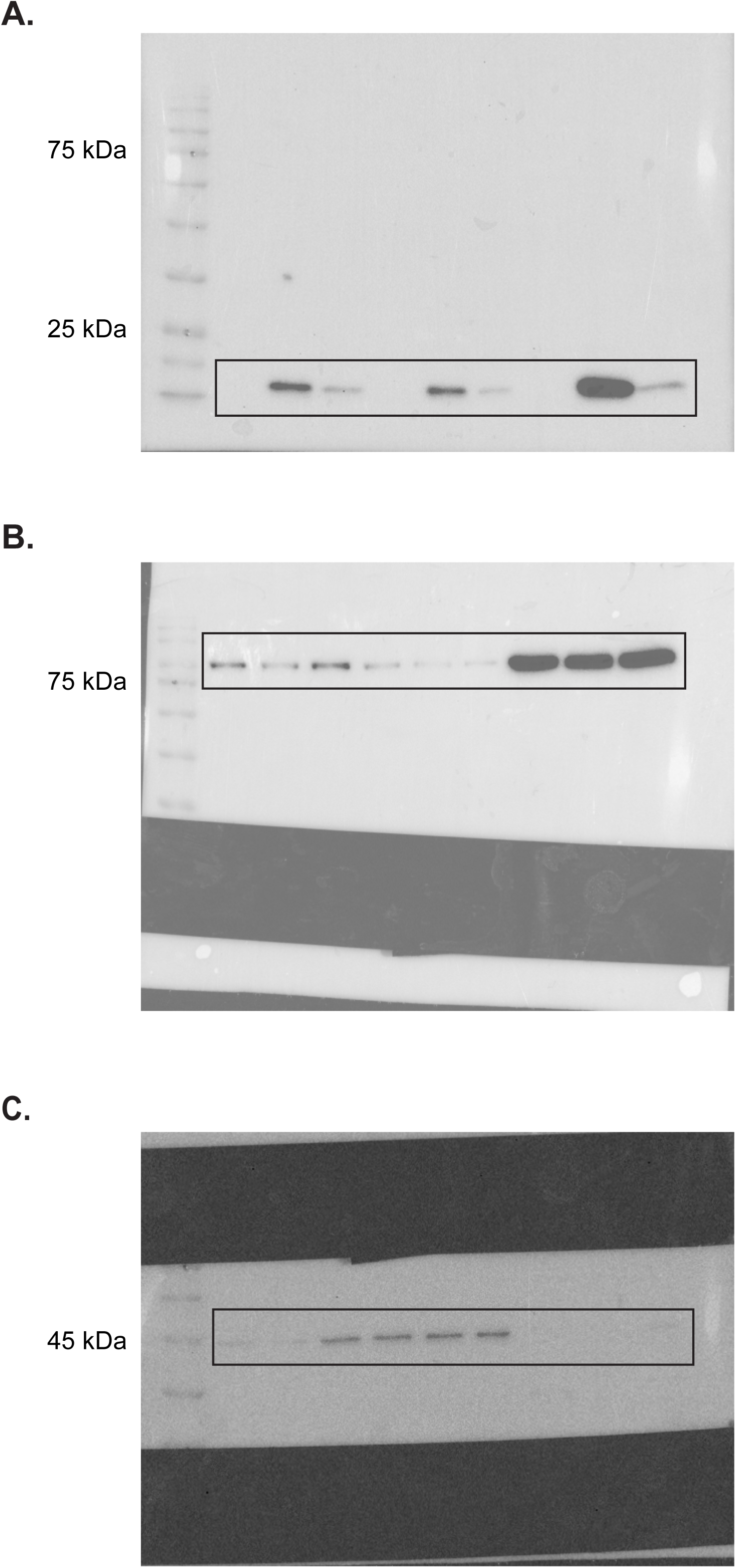

**Figure S6.**
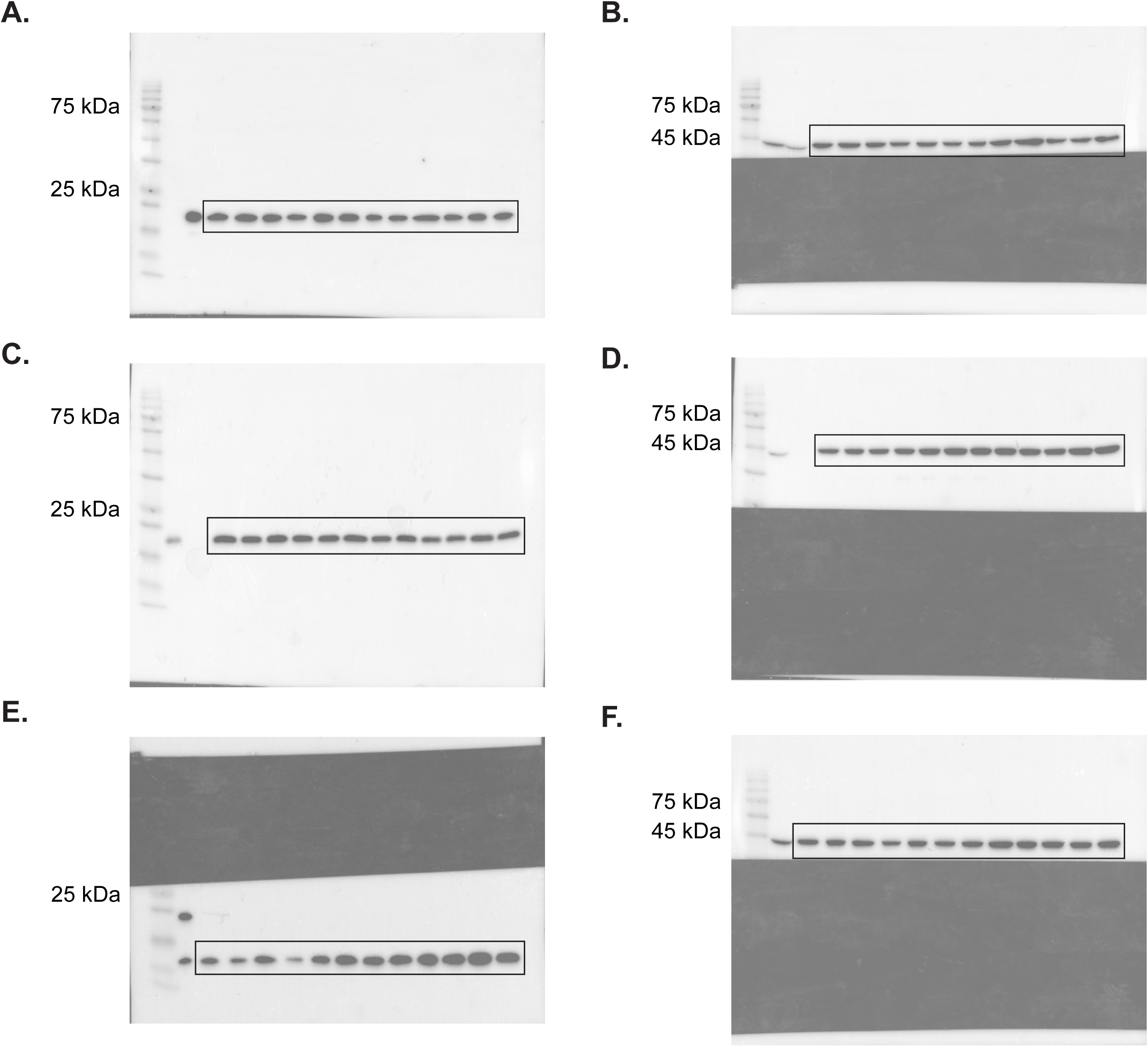

