## Supplementary Material for "A Kidney Stone–Associated CLDN4 Variant Impairs Tight Junction Stability and Paracellular Ion Permeability"

### 3. Protein Extraction and Immunoblotting

M1 cells were grown to confluence in 6-well plates and cultured under −DOX or +DOX conditions as indicated. Cells were lysed in RIPA buffer supplemented with protease inhibitors (Roche Complete Mini EDTA-free; cat. no. 04693159001) and phosphatase inhibitors (Roche PhosSTOP; cat. no. 04906837001). Lysates were clarified by centrifugation, and protein concentration was determined using a bicinchoninic acid (BCA) assay (Pierce, Thermo Fisher Scientific; cat. no. PI23225). Equal amounts of protein were resolved on 10 % or 12 % SDS–PAGE gels and transferred to polyvinylidene fluoride (PVDF) membranes. Membranes were blocked in 3 % nonfat dry milk prepared in Tris-buffered saline containing 0.1 % Tween-20 (TBST) and incubated overnight at 4 °C with primary antibodies. After incubation with horseradish peroxidase (HRP)–conjugated secondary antibodies, immunoreactive bands were detected using Immobilon Crescendo Western HRP substrate (MilliporeSigma; cat. no. WBLUR0500) and imaged with a ChemiDoc Touch imaging system (Bio-Rad). Densitometric analysis was performed using ImageJ (version 1.54g; National Institutes of Health). β-actin was used as the loading control for immunoblot analyses. Antibodies and dilutions are listed in **Table S1**.

4. **Cycloheximide (CHX) Chase Assay**

M1 cells expressing DOX-inducible HA-tagged WT or P74L CLDN4 were grown to confluence and induced with DOX (1 μg/mL for 24 h). To assess CLDN4 protein stability, culture medium containing DOX was aspirated at each time point and replaced with medium containing cycloheximide (MilliporeSigma; cat. no. 01810), prepared as a 10 mg/mL stock solution and used at a final concentration of 10 μg/mL to inhibit de novo protein synthesis. Cells were harvested at 0, 1, 2, 3, 4, 6, and 24 h following CHX addition, with all samples lysed at a common endpoint by adding CHX to the longest time point first and the 0 h time point last. Cell lysis, protein quantification, SDS–PAGE, immunoblotting, detection, and densitometric analysis were performed as described above. HA–CLDN4 band intensities were normalized to β-actin and expressed relative to the 0 h value for each biological replicate. Protein decay curves were fitted using a one-phase exponential decay model, and the half-life ($t_{1/2}$) was defined as the time at which normalized protein abundance reached 50 % of the initial level.

5. **Immunofluorescence Microscopy**

M1 cells expressing DOX-inducible HA-tagged WT or P74L CLDN4 were seeded on Transwell permeable filter supports (Corning; cat. no. 3401) and grown to confluence to allow epithelial polarization. Cells were then cultured in the presence or absence of DOX (1 µg/mL for 24 h) prior to immunofluorescence analysis. Cells were subsequently fixed with 4 % paraformaldehyde, permeabilized with 1 % Triton X-100, and blocked in 5 % bovine serum albumin. Sequential immunostaining was performed to assess CLDN4 localization relative to tight junction (TJs). Cells were first incubated with primary antibodies against the HA epitope, followed by the appropriate fluorophore-conjugated secondary antibodies. Antibodies and dilutions are listed in **Table S1**. Samples were subsequently incubated with primary antibodies against ZO-1, followed by corresponding secondary antibodies. Nuclei were counterstained with DAPI (MilliporeSigma; cat. no. D9564). Immunofluorescent imaging was performed using a WaveFX confocal microscope (Quorum Technologies, Guelph, Ontario, Canada) controlled by Volocity software (Quorum Technologies). Images were acquired with a 40× oil immersion objective, and Z-stack images were collected at 0.4-µm intervals. Images were processed uniformly for presentation.

### 6. Cell-Surface Biotinylation

M1 cells expressing DOX-inducible HA-tagged WT or P74L CLDN4 were seeded on Transwell permeable filter supports (Corning; cat. no. 3412) and grown to confluence until polarized. Cells were cultured under −DOX or +DOX conditions as indicated prior to cell-surface biotinylation. Cell-surface proteins were labeled using sulfo-N-hydroxysuccinimide–SS-biotin (Thermo Fisher Scientific; cat. no. 21331) essentially as described previously (2,3). Briefly, cells were washed on both apical and basolateral surfaces with ice-cold phosphate-buffered saline (PBS) and incubated with sulfo-NHS–SS-biotin (1.5 mg/mL in PBS) for 1 h at 4 °C. Excess biotin was quenched by washing with PBS containing 100 mM glycine, followed by a final wash in PBS. Cells were harvested and lysed in ice-cold RIPA lysis buffer supplemented with protease and phosphatase inhibitors as described above. Lysates were clarified by centrifugation, and protein concentration was determined using a BCA assay. Equal amounts of total protein were incubated with streptavidin–agarose beads (Thermo Fisher Scientific; cat. no. 53116) for 1 h at 4 °C with rotation. Following centrifugation, the supernatant was collected as the unbound (intracellular) fraction, and bead-bound proteins were eluted in Laemmli sample buffer to obtain the biotinylated (cell-surface) fraction. Input, unbound, and biotinylated fractions were analyzed by SDS–PAGE and immunoblotting. Imaging and densitometric analysis were performed as described above. For quantitative analysis of surface HA–hCLDN4, band intensities in the biotinylated fraction were normalized to the corresponding input for each biological replicate and expressed relative to the WT +DOX condition analyzed on the same membrane. $Na^+/K^+$-ATPase and β-actin were used as surface and cytosolic controls, respectively.

### 7. RNA Isolation and Quantitative Reverse Transcription Polymerase Chain Reaction (qRT-PCR)

Total RNA was extracted from M1 cells expressing WT or P74L CLDN4, cultured under induced (+DOX) or uninduced (–DOX) conditions, using TRIzol reagent (Invitrogen, Thermo Fisher Scientific; Cat. no. 15596018) according to the manufacturer's instructions. RNA yield and purity were determined using a NanoDrop™ 2000C spectrophotometer (Thermo Fisher Scientific). Reverse transcription was performed using 2 µg of total RNA with the High-Capacity cDNA Reverse Transcription Kit (Thermo Fisher Scientific; cat. no. 4368814) following the manufacturer's protocol. Quantitative real-time PCR was performed in technical triplicate for each cDNA sample using TaqMan PCR Master Mix on a QuantStudio 6 Pro qRT-PCR system (Thermo

Fisher Scientific). Gene expression was analyzed using the $2^{-\Delta\Delta CT}$ method (4). mRNA levels of target genes were normalized to the housekeeping gene ribosomal protein lateral stalk subunit P0 (*Rplp0*), whose expression was verified to remain stable across experimental conditions. Primer and probe sequences for each gene are listed in **Table S2**. For transcripts with low baseline expression, amplification yielded high Cq values; however, Cq distributions were comparable across cell lines and treatment conditions. Occasional undetermined values were assigned a Cq of 40 for ΔΔCT analysis.

**Supplementary Figure Legends**

**Supplementary Figure 1. qRT-PCR analysis of calcium transport–related genes in WT and P74L CLDN4–expressing M1 cells.** (A–E) Relative mRNA expression of *Trpv5*, *Trpv6*, *Calb1*, *Atp2b1*, and *Slc8a1* in WT and P74L CLDN4–expressing M1 cells under −DOX and +DOX conditions. Gene expression levels were normalized to *Rplp0* and expressed relative to the corresponding −DOX baseline for each cell line using the $2^{-\Delta\Delta CT}$ method. Data represent mean ± SEM from n = 3 biological replicates per condition. Statistical comparisons were performed using two-way ANOVA with Tukey's multiple comparisons test; no significant cell line– or DOX-dependent differences were detected. ns, not significant.

**Supplementary Figure 2. qRT–PCR analysis of γ-ENaC and ROMK in WT and P74L CLDN4–expressing M1 cells.** Relative mRNA expression of Scnn1g (γ-ENaC) (A) and Kcnj1 (ROMK) (B) in WT and P74L CLDN4–expressing M1 cells under −DOX and +DOX conditions. Gene expression levels were normalized to *Rplp0* and expressed relative to the corresponding −Dox baseline for each cell line using the $2^{-\Delta\Delta CT}$ method. Data represent mean ± SEM from n = 3 biological replicates per condition. Statistical comparisons were performed using two-way ANOVA with Tukey's multiple comparisons test; no significant cell line– or DOX-dependent differences were detected. ns, not significant.

**Supplementary Figure 3. Full-length immunoblots corresponding to Figure 2.** Uncropped membranes showing (A) HA–hCLDN4, (B) corresponding β-actin blot, (C) endogenous and human CLDN4, and (D) corresponding β-actin blot. Lanes (left to right) contain the molecular-weight ladder, IMCD kAE1 +DOX (HA-tag positive control), IMCD empty vector (EV) +DOX negative ontrol, and three biological replicates each of WT +DOX, WT −DOX, P74L +DOX, and P74L −DOX. Molecular-weight bands visible in each panel are indicated on the left (Panels A and C: 75 and 25 kDa; Panel B: 75 and 45 kDa; Panel D: 45 kDa). Boxes delineate the regions used for the cropped blots in Figure 2.

**Supplementary Figure 4. Full-length immunoblots corresponding to Figure 3.** Uncropped membranes showing the CHX chase for (A) WT HA–hCLDN4, (B) WT β-actin, (C) P74L HA–hCLDN4, and (D) P74L β-actin. Lanes (left to right) contain the molecular-weight ladder, IMCD kAE1 +DOX (HA-tag control), and samples collected at 0, 1, 2, 3, 4, 6, and 24 hours following CHX treatment. Visible molecular-weight markers are indicated on the left (Panels A and C: 25 kDa; Panels B and D: 45 kDa). Boxes delineate the regions used for the cropped blots in Figure 3.

**Supplementary Figure 5. Full-length immunoblots corresponding to Figure 4B.** Uncropped membranes showing (A) HA–hCLDN4, (B) $Na^+/K^+$-ATPase, and (C) β-actin from the cell-surface biotinylation experiment. Lanes (left to right) contain the molecular-weight ladder, followed by Input, Unbound, and Biotinylated fractions for WT −DOX, WT +DOX, and P74L +DOX cells. Visible molecular-weight markers are indicated on the left (Panel A: 75 and 25 kDa; Panel B: 75

kDa; Panel C: 45 kDa). Boxes delineate the regions used for the cropped blots shown in Figure 4B.

**Supplementary Figure 6. Full-length immunoblots corresponding to Figure 5.** Uncropped membranes showing tight-junction protein expression in WT and P74L CLDN4–expressing M1 cells under +DOX and −DOX conditions. Panels display full blots for (A) CLDN3, (B) its corresponding β-actin, (C) CLDN7, (D) corresponding β-actin, (E) CLDN8, and (F) corresponding β-actin. CLDN3 blots: Lanes (left to right) contain the molecular-weight ladder, kidney lysate, liver lysate, and three biological replicates each of WT +DOX, WT −DOX, P74L +DOX, and P74L −DOX. Visible molecular-weight markers are indicated on the left (Panel A: 75 and 25 kDa; Panel B: 75 and 45 kDa). CLDN7 blots: Lanes include the molecular-weight ladder, kidney lysate, an empty lane, and three biological replicates each of WT +DOX, WT −DOX, P74L +DOX, and P74L −DOX. Molecular-weight markers are shown at 75 and 25 kDa (Panel C) and 75 and 45 kDa (Panel D). CLDN8 blots: Lanes contain kidney lysate followed by three biological replicates each of WT +DOX, WT −DOX, P74L +DOX, and P74L −DOX. Molecular-weight markers are shown at 25 kDa (Panel E) and 75 and 45 kDa (Panel F). Boxes indicate the cropped regions corresponding to blots in Figure 5A, 5C, and 5E.

### Supplementary Tables

#### Table S1. Antibodies used in this study

| Target | Application | Host | Type | Conjugate | Supplier | Cat. no. | Dilution |
|---|---|---|---|---|---|---|---|
| **Primary antibodies** | | | | | | | |
| CLDN3 | WB | Rabbit | Primary | — | Invitrogen | 34-1700 | 1:1000 |
| CLDN4 | WB | Rabbit | Primary | — | Invitrogen | 36-4800 | 1:500 |
| CLDN7 | WB | Rabbit | Primary | — | Invitrogen | 34-9100 | 1:1000 |
| CLDN8 | WB | Rabbit | Primary | — | Invitrogen | 40-0700Z | 1:1000 |
| HA | IF | Mouse | Primary | — | BioLegend | 901513 | 1:500 |
| HA | WB | Mouse | Primary | — | Cell Signaling Technology | 2367 | 1:1000 |
| $Na^+/K^+$-ATPase | WB | Mouse | Primary | — | Santa Cruz Biotechnology | sc-48345 | 1:5000 |
| ZO-1 | IF | Rat | Primary | — | DSHB | R26.4C-s | 1:25 |
| **Loading control** | | | | | | | |
| β-actin | WB | Mouse | Primary | HRP | BioLegend | 643807 | 1:10000 |
| **Secondary antibodies** | | | | | | | |
| Mouse IgG | IF | Donkey | Secondary | Cy3 | Jackson ImmunoResearch | 715-165-151 | 1:500 |
| Mouse IgG | WB | Horse | Secondary | HRP | Cell Signaling Technology | 7076S | 1:10000 |
| Rabbit IgG | WB | Goat | Secondary | HRP | Cell Signaling Technology | 7074P2 | 1:10000 |
| Rat IgG | IF | Donkey | Secondary | Alexa Fluor 488 | Thermo Fisher Scientific | A-21208 | 1:500 |

**Table S2. Primer and probe sequences of mouse genes used in qRT-PCR analysis**

| Gene | Assay Name | Forward Primer | Reverse Primer | Probe |
|---|---|---|---|---|
| *Atp2b1* | Mm.PT.58.12757227 | 5′-CTCACTGGCTTACTCAGTCAAG-3′ | 5′-CGTCATCCTGTTCATCGTCA-3′ | 5′-/56-FAM/AGCTGTAGC/ZEN/ATTCCCCATGGTCTC/3IABkFQ/-3′ |
| *Calb1* | Mm.PT.58.23638207 | 5′-CTGATCACAGCCTCACAGTT-3′ | 5′-TTTCCGGTGATAGCTCCAATC-3′ | 5′-/56-FAM/AACCACTTC/ZEN/CGTCAGCGTCGAA/3IABkFQ/-3′ |
| *Cldn4* | N009903.1.pt.mCldn4 | 5′-CAGGACTGCCAAAGGAGATTC-3′ | 5′-AACACTTTCTCAGCCCTCTG-3′ | 5′-/56-FAM/CATAGACGC/ZEN/CATCGCTCAGCCTC/3IABkFQ/-3′ |
| *Cldn8* | N018778.1.pt.mCldn8 | 5′-CCACTGAGGCATGATAGTCAC-3′ | 5′-AGCTGGATACAATTTGGGAGG-3′ | 5′-/56-FAM/TGCAGCCAT/ZEN/TTGAAGAGCGTAGGT/3IABkFQ/-3′ |
| *Kcnj1* | NM_001168354.1 | 5′-CAGGACTTCGAGTTGGTTGT-3′ | 5′-GCACCTCTTCTGGGATGTATG-3′ | 5′-/56-FAM/TAGAATCCA/ZEN/CCAGTGCAACCTGCC/3IABkFQ/-3′ |
| *Rplp0* | Mm.PT.58.43894205 | 5′-TTATAACCCTGAAGTGCTCGA-3′ | 5′-CGCTTGTACCCATTGATGATG-3′ | 5′-/56-FAM/AGGCCTGC/ZEN/ACTCTCGCTT/3IABkFQ/-3′ |
| *Scnn1g* | N011326.1.pt.scnn1g | 5′-GTCAGAGGTGTCATTTGAGCA-3′ | 5′-GAGAACGAGAAGGGAAAGGC-3′ | 5′-/56-FAM/TCGGAAGCG/ZEN/GAAAATCAGGGGAA/3IABkFQ/-3′ |
| *Slc8a1* | Mm.PT.58.9647961 | 5′-TTTCTCATCCAGCCCATGC-3′ | 5′-CCAGACGAAATCCCATTGAAAC-3′ | 5′-/56-FAM/CTTCCAACT/ZEN/GCTCCAACCTCTGTCA/3IABkFQ/-3′ |
| *Trpv5* | Mm.PT.58.11633143 | 5′-ACAGAAATCACTACAGGAAGCA-3′ | 5′-GCACCAACTCTGAAGATGTCT-3′ | 5′-/56-FAM/AGCCCCAAT/ZEN/GACTGTCACCAACT/3IABkFQ/-3′ |
| *Trpv6* | Mm.PT.58.42025701.g | 5′-GTGGGTAGTGAGGAGATTGTTAG-3′ | 5′-TTTGTTGGGCTGCAGGAT-3′ | 5′-/56-FAM/ACGGATGTC/ZEN/AGCTCCATGCTCAAT/3IABkFQ/-3′ |

**Table S3. Ussing Chamber Solutions**

| Component (mM) | Control | Low $Na^+$ | Control – no phosphate | High $Ca^{2+}$ |
|---|---|---|---|---|
| NaCl | 143.6 | 30 | 140 | 0 |
| KCl | 0 | 0 | 3.6 | 3.6 |
| $KH_2PO_4$ | 0.4 | 0.4 | 0 | 0 |
| $K_2HPO_4$ | 1.6 | 1.6 | 0 | 0 |
| $MgCl_2$ | 1 | 1 | 1 | 1 |
| $CaCl_2$ | 0 | 0 | 0 | 70 |
| Calcium gluconate | 1.3 | 1.3 | 1.3 | 0 |
| HEPES | 0 | 0 | 3 | 3 |
| Glucose | 10 | 10 | 10 | 10 |
| Mannitol | 0 | 227 | 5 | 78 |
